# Targeting the Hedgehog Signaling Pathway to Improve Tendon-to-Bone Integration

**DOI:** 10.1101/2022.09.27.509810

**Authors:** Timur B. Kamalitdinov, Keitaro Fujino, Sinaia Keith Lang, Xi Jiang, Rashad Madi, Mary Kate Evans, Miltiadis H. Zgonis, Andrew F. Kuntz, Nathaniel A. Dyment

**Affiliations:** Department of Orthopaedic Surgery, University of Pennsylvania, Philadelphia, PA; Department of Bioengineering, University of Pennsylvania, Philadelphia, PA; Osaka Medical College, Osaka Prefecture, Takatsuki, Japan

**Keywords:** hedgehog signaling, ACL reconstruction, tendon-to-bone repair, mineralized fibrocartilage, transgenic mice

## Abstract

**Objective:** While the role of Hedgehog (Hh) signaling in promoting zonal fibrocartilage production during development is well-established, whether this pathway can be leveraged to improve tendon-to-bone repair in adults is unknown. Our objective was to genetically and pharmacologically stimulate the Hh pathway in cells that give rise to zonal fibrocartilaginous attachments to promote tendon-to-bone integration.

**Design:** Hh signaling was stimulated genetically, via constitutive Smo (SmoM2 construct) activation of activated bone marrow stromal cells, or pharmacologically, via systemic agonist delivery, to mice following anterior cruciate ligament reconstruction (ACLR). To assess tunnel integration, we measured mineralized fibrocartilage (MFC) formation in these mice 28 days post-surgery and performed tunnel pullout testing.

**Results:** Hh pathway-related genes increased in cells forming the zonal attachments in WT mice. Both genetic and pharmacologic stimulation of the Hh pathway increased MFC formation and integration strength 28 days post-surgery. We next conducted studies to define the role of Hh in specific stages of the tunnel integration process. We found Hh agonist treatment increased proliferation of the progenitor pool in the first week post-surgery. Additionally, genetic stimulation led to continued MFC production in later stages of the integration process. These results indicate that Hh signaling plays an important biphasic role in cell proliferation and differentiation towards fibrochondrocytes following ACLR.

**Conclusion:** This study reveals a biphasic role for Hh signaling during the tendon-to-bone integration process after ACLR. In addition, the Hh pathway is a promising therapeutic target to improve tendon-to-bone repair outcomes.

## INTRODUCTION

Approximately 30% of adults in the United States suffer from a musculoskeletal condition, with an increasingly aging population leading to an increased prevalence of tendon and ligament injuries^1^. Many of these injuries occur at tendon-to-bone insertion sites, or entheses. Entheses contain four distinct tissue zones beginning with the tendon midsubstance which transitions into unmineralized fibrocartilage followed by mineralized fibrocartilage and ultimately the underlying bone. These interfaces are critical for tendon function as they facilitate load transfer between the stiff bone and relatively compliant tendon^2,3^. The stiffness mismatch between tendon and bone in the enthesis yields stress concentrations, contributing to the high prevalence of injuries at the enthesis that do not spontaneously heal^4–6^. Traditional tendon-to-bone reattachment strategies where the tendon is anchored to the bone with sutures result in disorganized scar formation instead of reestablishing the zonal architecture, leading to impaired load transfer and significant re-tear rates^7,8^. Therefore, there is a critical need to improve current tendon-to-bone repair strategies. Unlike traditional tendon-to-bone repair, ligament reconstructions where tendon grafts are passed through bone tunnels can produce zonal attachments that share key characteristics with entheses such as collagen fibers traversing unmineralized and mineralized fibrocartilage zones^9–19^. This model system provides a platform to investigate mechanisms that regulate tendon-to-bone integration following injury.

During normal tendon growth and development, there is a coordinated series of events that lead to zonal enthesis formation^20^. Beginning with the specification of the enthesis progenitor pool^20–24^, these cells expand and proliferate as they synthesize the collagenous matrix. A subset of these cells differentiates into fibrochondrocytes that deposit proteoglycan-rich matrix to create the unmineralized fibrocartilage zone. Finally, a portion of unmineralized fibrochondrocytes closest to the bone interface mineralize the surrounding matrix to create the mineralized fibrocartilage zone^20^. Similarly, adult anterior cruciate ligament (ACL) reconstructions produce zonal tendon-to-bone attachments between the tendon graft and adjacent newly formed bone through a staged repair response. Using a murine ACL reconstruction (ACLR) model, we previously demonstrated that an alpha smooth muscle actin (αSMA)-expressing amplifying progenitor pool of bone marrow stromal cells (bMSCs) adjacent to the inserted tendon graft, not the tenocytes within the graft, initiate the zonal attachment formation process by infiltrating the graft and anchoring collagen fibers to the adjacent bone^18,19^. The cells in these nascent attachment sites then synthesize proteoglycan-rich fibrocartilage and ultimately mineralize this fibrocartilage to form four distinct zones within the tendon-to-bone attachment. However, the cell signaling events responsible for this staged repair response are not clear.

One such cell signaling pathway that is critical to enthesis development and may play a similar role in tendon-to-bone repair in the adult is the hedgehog pathway. During embryonic development, the downstream hedgehog (Hh) transcription factor Gli1 is one of the distinct markers that delineates enthesis progenitors from adjacent tendon midsubstance or nascent cartilage cells. These Gli1-expressing progenitors give rise to the unmineralized and mineralized fibrochondrocytes of the fully formed zonal enthesis^23^. Hh signaling is a potent positive regulator of fibrocartilage differentiation and maturation^20,23–27^ as Hh overactivation in Scx-expressing tendon cells leads to ectopic fibrocartilage formation in the tendon midsubstance^24^ while ablating Hh signaling leads to striking deficits in enthesis mineralized fibrocartilage (MFC) formation^20,23,25^. Furthermore, Hh signaling ligands play a biphasic role in enthesis development^20,23–25,27^ with Sonic hedgehog (Shh) regulating the specification of enthesis progenitors during embryonic development^27^ and Indian hedgehog (Ihh) promoting enthesis maturation during postnatal growth^20,24^. Therefore, Hh signaling plays critical roles in promoting zonal enthesis fibrocartilage specification, differentiation, and maturation.

Since Hh signaling is a potent regulator of enthesis fibrocartilage formation during growth and development, we hypothesize that it plays a similar role in fibrocartilage production during tendon-to-bone repair. In fact, the pathway is active during healing after tendon-to-bone repair of the rotator cuff in rats^28^ and ACL in rats and mice^13,29,30^. However, it is not clear what role the pathway has in regulating the repair response after surgery. Therefore, the objective of our current study is to genetically and pharmacologically stimulate the Hh signaling pathway to promote tendon-to-bone integration following ACLR. We utilize conditional Hh overexpression transgenic mice or systemic Hh agonist injections to stimulate the Hh pathway in cells that contribute to the repair response. The extent of zonal tunnel integration is assessed using multiplexed mineralized cryohistology and tunnel pullout tests. An improved understanding of the signaling pathways regulating tendon-to-bone attachment formation during repair is a crucial step towards developing new therapies to improve repair outcomes.

## METHODS

### Mice

All animal procedures were approved by the University of Pennsylvania’s Institutional Animal Care and Use Committee. The transgenic mouse lines used in this study were described previously: αSMACreERT2^31^, R26R-tdTomato Cre reporter (B6;129S6-*Gt(ROSA)26Sor*^*tm9(CAG-tdTomato)Hze*^/J, stock # 007905)^32^, and R26SmoM2 (STOCK *Gt(ROSA)26Sor*^*tm1(Smo/EYFP)Amc*^/J, stock # 005130)^33^. αSMACreERT2 and R26R-tdTomato mice were bred in a mixed genetic background containing C57BL/6, CD1, and 129/SvJ strains. R26SmoM2 mice were maintained on a C57BL/6 background. αSMACreERT2 mice were crossed with R26SmoM2 mice or R26R-tdTomato mice to yield double transgenic mice.

### Experimental design

ACL reconstructions were performed on a total of 71 mice (mean ± SD age, 16.5 ± 2.3 weeks old). To measure expression of Hh-and enthesis-related genes, αSMACreERT2;R26R-tdTomato mice were sacrificed at 3, 7, and 14 days post-surgery for qPCR analysis from microdissected histological sections (n = 3-4/timepoint). To increase Hh activity genetically in cells that give rise to the tendon-to-bone attachments, αSMACreERT2;R26SmoM2 (SmoCA) mice received intraperitoneal tamoxifen injections (80 mg/kg) (Sigma-Aldrich Corp.) on the day of surgery and every other day thereafter for a total of five injections to constitutively activate Smo in αSMA-expressing cells. Demeclocycline (Sigma-Aldrich Corp.) injections (60 mg/kg) were given the day prior to sacrifice to label deposited mineral. A subset of mice also received calcein (6 mg/kg) (Sigma-Aldrich Corp.) seven days prior to sacrifice to label deposited mineral at that time. Mice were sacrificed at 28 days post-surgery and assigned to cryohistology (n = 10-13/group) or biomechanics (n = 5-7/group). To increase Hh activity pharmacologically, αSMACreERT2;R26R-tdTomato mice received either the Hh agonist (Hh-Ag1.5, Cellagen Technology, 20 mg/kg) or PBS injections five times per week. These mice were sacrificed at 28 days post-surgery and assigned to cryohistology (n = 6-7/group) or biomechanics (n = 7/group). Additional agonist-treated and PBS control mice were assigned to proliferation and microdissected qPCR analysis at day 7 post-surgery. These mice received tamoxifen injections on the day of surgery, 3-, and 5-days post-surgery along with 5-Ethynyl-2′-deoxyuridine (EdU, Click Chemistry Tools, 3 mg/kg) every day (n = 4-5/group). Cryosections were either collected for qPCR analysis or stained for EdU. Both male and female sexes were included in this study and were equally distributed across treatment groups.

### ACL reconstruction procedure using tail tendon autografts

The right knee joint of each mouse was subjected to surgical transection of the anterior cruciate ligament followed by reconstruction using tail tendon autografts, similar to previously described work^18,19^. Mice were anesthetized with isoflurane (1-3%) given pre-surgical analgesia, and sterilely prepped. To acquire the graft tissue, 8-10 tail tendon fascicles (length = 3-4 cm) were harvested from the same mouse as a bundle and maintained in PBS to prevent dehydration. An anteromedial incision was made adjacent to the patellar tendon and the patella was subluxed laterally to access the joint space. The ACL was transected with a 27G needle at the femoral insertion. After confirmation of substantial anterior drawer and an intact posterior cruciate ligament, a 27G needle was used to hand drill the tibial tunnel originating as close to the native ACL footprint as possible and exiting on the anteromedial cortex of the tibia within the metaphysis. Next, a new 27G needle was used to hand drill the femoral tunnel originating at the native ACL footprint and exiting on the lateral surface of the femur proximal to the patella. A 27G needle was inserted antegrade into the tibial tunnel and suture was wrapped around the mid-length of the tail tendon bundle and passed through the needle. Once the suture, but not the tail tendon bundle, was through the needle, the needle was removed, and the tendon graft was pulled through the tunnel. A similar procedure was performed to pass the tendon graft through the femoral tunnel. Once the mid-length of the tendon graft was outside of the femoral tunnel, it was passed through and around a 316 stainless-steel washer (OD 1.98mm, ID 1.0 mm; McMaster Carr) to anchor the graft to the washer (i.e., cow hitch knot) at the outer cortex of the femur. The knee was positioned near full extension and then the two ends of the tail tendon graft at the outer cortex of the tibia were tied to another stainless-steel washer using a surgical knot such that the washer was positioned as close to the outer cortex as possible. The patella was then placed back to its anatomical position and the patellar tendon and medial retinaculum were sutured closed followed by skin closure. After recovery from anesthesia, the animals were returned to their cage and allowed to move ad libitum. At assigned timepoints, mice were euthanized by CO_2_ asphyxiation.

### qPCR of microdissected sections

For gene expression analysis at different stages of the repair process after ACLR, mice were euthanized at days 3, 7, and 14 post-surgery. Hindlimbs were harvested and quickly transferred to a 4% phosphate-buffered paraformaldehyde solution and were fixed for 4 hours on a shaker at 4°C. The fixed samples were embedded in optimal cutting temperature compound (OCT, Scigen) and kept at -80°C to minimize RNA degradation. Tape-stabilized 20µm thick tissue sections were collected and kept frozen until tissue capture. Specific regions from each timepoint were collected by cutting the tissue and underlying tape with the bevel of a 27G needle and were transferred to a blank piece of cryofilm to pool pieces of tissue from multiple sections per sample (Fig. S1). Regions of the expanding marrow (defined as the fibrotic marrow adjacent to the tunnel interface), early attachments, and mineralized attachments were collected from the day 3, 7, and 14 samples, respectively. Native ACL entheses were collected from the contralateral limbs. Cryofilm pieces containing the tissues were submerged in digest solution consisting of 1X Digestion Buffer (Zymo Research) and 1 mg/ml Proteinase K (Zymo Research) for 1 hour at 55°C. Samples were further de-crosslinked at 65°C for 15 minutes. After digestion, RNA isolation was performed using the Quick-RNA MicroPrep kit (Zymo Research) according to the manufacturer’s protocol and as previously described^34^. For cDNA synthesis, the SuperScript IV VILO kit was used (Thermo Fisher Scientific) following ezDNAse treatment to remove gDNA. To pre-amplify the cDNA for the specific gene targets prior to qPCR, a primer pool of the 20X Taqman probes was first generated and diluted 1:100 in DNA suspension buffer (1X TE buffer). Then, the Fluidigm Preamp Master Mix was used to pre-amplify the cDNA for specific gene targets for 15 cycles and the resultant cDNA product was diluted 1:5 with DNA suspension buffer. qPCR was performed on the pre-amplified cDNA samples using the Taqman Fast Advanced Master Mix (Thermo Fisher Scientific). Taqman probes for 18S, Rps17 (housekeepers), Acta2, Col1a1, Col2a1, Col10a1, Ptch1, Smo, Gli1, and Ihh (Taqman IDs in order: Mm03928990_g1, Mm01314921_g1, Mm00725412_s1, Mm00801666_g1, Mm01309565_m1, Mm00487041_m1, Mm00436026_m1, Mm01162710_m1, Mm00494654_m1, Mm00439613_m1) were used. Principal component analysis (PCA) and hierarchical clustering was performed on ΔC_T_ values using ClustVis^35^.

For gene expression analysis of the subset of mice that received Hh agonist or vehicle injections for 7 days, mice were euthanized at 7 days post-surgery. Regions of expanding marrow around both sides of the femoral tunnel were captured and used for qPCR analysis with Taqman probes for 18S, Gli1, and Ptch1 as described above.

### Multiplexed mineralized cryohistology

Following euthanasia at assigned timepoints, hindlimbs were harvested and fixed in formalin for 3 days, transferred to 30% sucrose overnight, and embedded in OCT. Tape-stabilized 8µm thick frozen mineralized sagittal sections^36^ of the knee were collected and each section was subjected to multiple rounds of imaging on the Zeiss Axio Scan.Z1 digital slide scanner. For samples assigned to histology at 28 days after surgery, imaging rounds included 1) fluorescent reporters, mineralization labels, and polarized light, 2) alkaline phosphatase (AP) fluorescent staining (ELF97 Phosphatase Substrate, Thermo Fisher Scientific) with Hoechst 33342 counterstain (Invitrogen), and 3) 0.025% toluidine blue (TB). Sections from mice assigned to histology 7 days after surgery underwent imaging rounds including 1) fluorescent reporters, polarized light, and Calcein Blue staining (Sigma-Aldrich Corp.)^36^, 2) EdU detection with Alexa Fluor 647 azide (Thermo Fisher Scientific) and Click-&-Go Cell Reaction Buffer Kit (Click Chemistry Tools), and 3) Hematoxylin and aqueous eosin staining (Thermo Fisher Scientific). Layered composite images of all imaging rounds were assembled and aligned in image editing software.

### Tunnel pullout test

Tunnel pullout tests were performed similarly to previous work^18^. Briefly, following euthanasia, mice assigned to biomechanics were frozen at -20°C until the day of testing. After thawing, limbs were dissected at the hip joint and the femur was potted in polymethyl methacrylate (PMMA). A 15-pound test braided fishing line (PowerPro) was passed through the washer on the outer cortex of the femur and the potted limb was mounted on a ball head tripod (Manfrotto) attached to a custom fixture. A custom grip fixture was attached to the Instron (Model 5542a) load cell on one end and the fishing line on the other end. After aligning the vertical length of the fishing line with the bone tunnel, the femur was loaded uniaxially at 0.025 mm/sec until failure. Maximum loads were recorded for pullout strength.

### Image analysis

The mineralized fibrocartilage area was assessed at 28 days post-surgery for each treatment group by identifying the regions within the tunnel containing highly aligned collagen fibers (polarized light) traversing a fluorescent tidemark (i.e., demeclocycline) and normalizing these areas by the length of the tunnel using Fiji (ImageJ)^37^.

The percentage of tunnel length containing MFC at 28 days post-surgery was quantified in Fiji by using the specific region of MFC area and tunnel length line for a particular section and performing the Plot Profile function to generate a two-dimensional graph of pixel intensities along the length of the line. The width of the line was set to encompass all MFC areas in the section, and the percentage of nonzero pixel intensities along the line were normalized by the total number of pixels from the plot profile to generate the percentage of tunnel length containing MFC for a particular section.

The percentage of MFC produced in the last week in mice at 28 days post-surgery was assessed in Fiji by overlaying the MFC areas quantified previously onto the original composite image for a particular section. Then, confined by these MFC boundaries of the mineral state at 28 days post-surgery, the calcein area, denoting MFC that was present at 21 days post-surgery at the time of calcein injection, was quantified. The difference between total MFC area and the calcein area represented the mineralized fibrocartilage produced in the timeframe between the two fluorescent mineral labels. This differential area was then normalized by the total MFC area of the section. The MFC areas before and after day 21 from this analysis were also normalized by the tunnel length.

To quantify EdU-positive cells in the expanding marrow around the tendon graft, a binary nuclear mask was created from the Hoechst 33342-stained imaging round in Fiji, the selection for analysis was drawn around the regions of expanding marrow, and the Analyze Particles function was used after redirecting the measurement to the EdU channel image. To exclude EdU-positive cells within newly formed woven bone, the Analyze Particles function was used again after redirecting to a corresponding binarized Calcein Blue channel image and cells that expressed positive EdU and Calcein Blue signal were excluded during further analysis.

Bone parameters were measured in mice at 28 days post-surgery using OsteoMeasure (OsteoMetrics). Sagittal sections of the femoral tunnel from MFC analysis were used. Regions to define total area were drawn approximately 400µm away from each side of the tunnel-bone interface. Bone area fraction (BAF) was then calculated by outlining the newly formed bone in these regions, excluding the growth plate, and dividing by the total area. Mineral apposition rate (MAR) was measured by tracing paired green (calcein) and yellow (demeclocycline) labels in the newly formed bone in the defined regions and calculated in OsteoMeasure. Mineralizing surface/bone surface (MS/BS) was measured by tracing demeclocycline-positive surfaces in the newly formed bone and dividing by the total bone perimeter. Similarly, alkaline phosphatase surface staining/bone surface (AP/BS) was traced along the newly formed bone surfaces and normalized by the total bone perimeter.

### Statistics

All statistical analyses were performed using GraphPad Prism 9.2.0 (GraphPad Software). Normally distributed data were confirmed using D’Agostino-Pearson and Shapiro-Wilk normality tests. Treated and control groups for their respective datasets were compared using Student’s t-tests (p < 0.05). Data are presented as mean ± SD.

## RESULTS

### Hh-related gene activity increases during tendon-to-bone integration

Following ACL reconstruction, tendon-to-bone integration occurs through a series of coordinated events^9–19,38–41^. We wanted to investigate the spatiotemporal expression pattern of Hh-related genes during these stages. Therefore, we captured specific tissue regions from fixed sections of tissue at days 3, 7, and 14 post-surgery. In the day 3 sections, we captured tissue from regions of the expanding marrow, which is characterized by areas of denser extracellular matrix (eosin in Fig. 1A) containing αSMA-expressing cells^18^ adjacent to the tendon graft. In day 7 sections, we captured tissue from regions where αSMA+ cells began to infiltrate the tendon graft and initiate the early stages of the attachment process (Fig. 1B). Finally, we captured tissue from regions of the mineralized attachment sites in day 14 sections, which could be visualized well with αSMACreERT2;R26R-tdTomato expression in areas of strong demeclocycline labeling (Fig. 1C). These attachment sites were more mature compared to those seen at day 7, which did not contain mineralized fibrocartilage. Contralateral ACL entheses were captured from both day 3 and day 7 samples as controls. Detailed methods and representative images of captured tissue regions for each timepoint are shown in Fig. S1.

**Figure 1.**
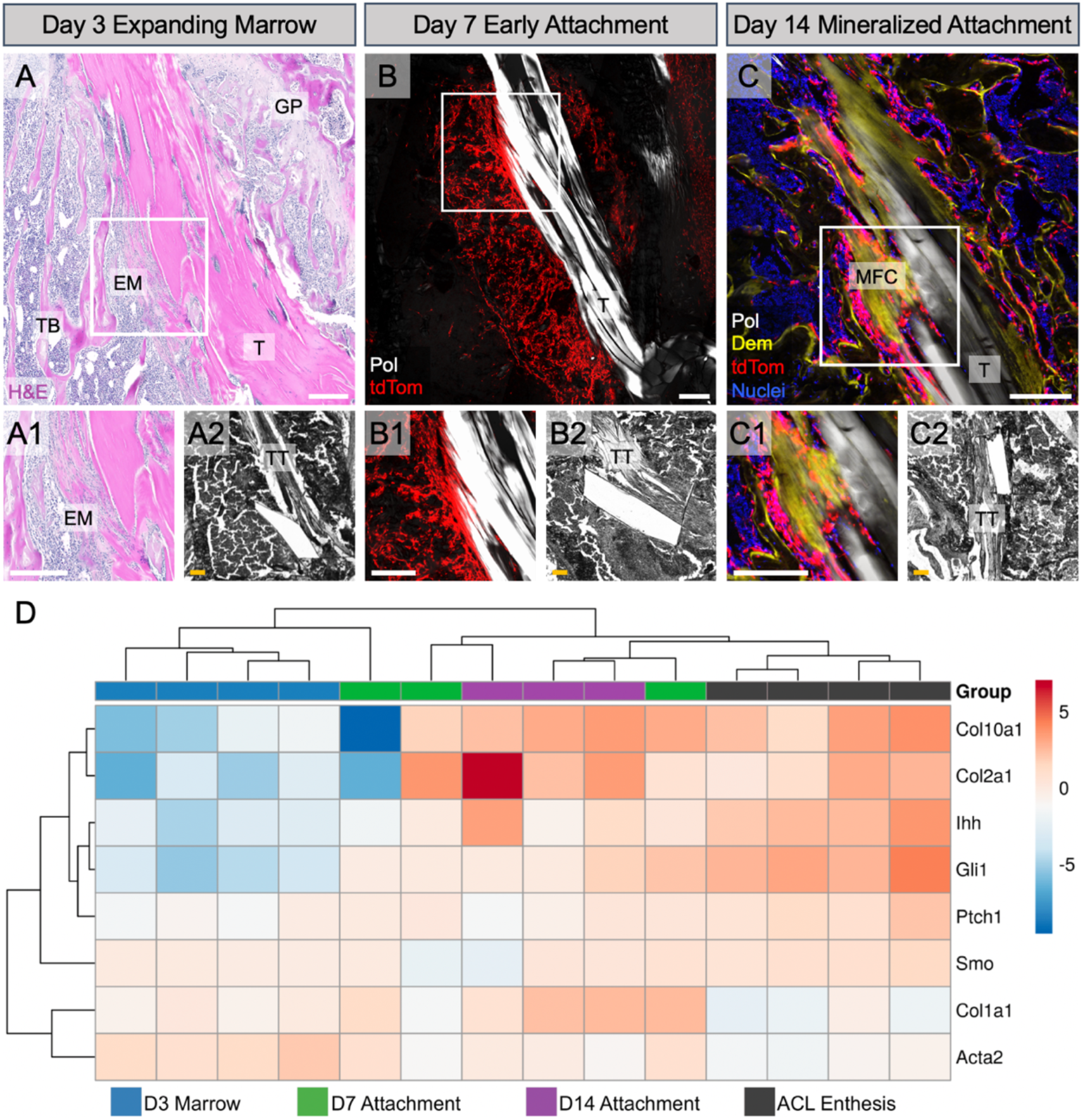
Hh-related gene activity increased during tendon-to-bone integration after ACLR. At day 3 post-surgery, bone marrow cells amplified and expanded around the tendon graft, resulting in expanding marrow (EM) with denser ECM indicated by stronger eosin staining (A, A1). At 7 days post-surgery, the amplified cells as visualized with the αSMACreERT2;R26R-tdTomato model began to infiltrate the tendon graft and form early attachment sites (B, B1). By day 14 post-surgery, there were mineralized attachments present containing mineralized fibrocartilage (MFC) where collagen fibers (Pol light, white) span an area of strong demeclocycline (Dem) signal (C, C1). We captured these specific regions at each timepoint for qPCR, where panels A-C are for demonstration purposes and A2, B2, and C2 demonstrate representative captured tissue regions at those timepoints. Contralateral ACL enthesis tissue was also collected. A qPCR (ΔC_T_) heatmap (D) was generated after analyzing captured tissues for fibrochondrocyte (Col10a1, Col2a1), Hh-related (Ihh, Gli1, Ptch1, Smo), and ECM (Col1a1, Acta2) genes (n = 3-4/group). T = tunnel, TT = tail tendon, GP = growth plate, TB = trabecular bone. Scale bars = 200μm.

We then queried Hh-related gene expression changes, in addition to other markers expressed by cells in these regions, throughout the integration process using qPCR. We performed principal component analysis with hierarchical clustering on ΔC_T_ values obtained from qPCR for all samples and target genes. Hierarchical clustering revealed that samples from each timepoint generally clustered together. As expected, cells from regions of expanding marrow (day 3) expressed high levels of Acta2 as the cells activated and expanded in response to the surgery, while Acta2 expression was lower in cells within the attachments in later stages of repair (Fig. 1D). These results correspond with previous IHC data^18^. Additionally, Col1a1 levels increased throughout the repair process as cells infiltrated the graft and deposited collagen matrix to anchor the newly formed attachments to bone, while cells in the contralateral native ACL enthesis had lower Col1a1 expression. As expected, the ACL entheses and day 14 mineralized attachments had higher expression of fibrocartilage markers Col2a1 and Col10a1, compared to the expanding marrow and early attachments. Finally, the Hh-responsive gene Gli1 and the ligand Ihh were highly expressed in ACL entheses and mineralized attachment sites, correlating with mature enthesis fibrocartilage markers, whereas Ptch1 and Smo were relatively consistent during the repair process. Overall, these expression data indicate that Hh activity increases in cells as they form mature tendon-to-bone attachments.

### Hh signaling promotes zonal attachment formation during tunnel integration following ACL reconstruction

To investigate the role of Hh signaling in regulating the repair response after surgery, we utilized both genetic and pharmacologic approaches to stimulate the pathway in cells participating in the repair response. In our genetic approach, we crossed αSMACreERT2 mice, which we previously showed target the amplifying bone marrow mesenchymal progenitors that contribute to zonal tendon-to-bone attachment formation^18^, with mice containing the R26SmoM2 construct harboring a W539L point mutation in the central signal transducer of the Hh pathway Smoothened (Smo) (Fig. 2A). Upon tamoxifen administration, these mice experienced constitutive Smo expression (SmoCA) in αSMA-expressing cells and their progeny. To assess the extent of zonal tunnel integration after ACL reconstruction, we used mineral labeling (demeclocycline) and collagen structure (polarized light) in histological sections to identify regions of mineralized fibrocartilage (MFC), which we then quantified and normalized to bone tunnel length (Fig. 2B). With increased Hh signaling, via Smo over-expression (SmoCA) in αSMA-lineage cells (Fig. 2D), the SmoCA mice had 36% greater MFC area than controls at 28 days post-surgery (p = 0.02, Fig. 2E, F). This result indicated that Hh signaling in these cells promotes the production of zonal tendon-to-bone attachments in the tunnels.

**Figure 2.**
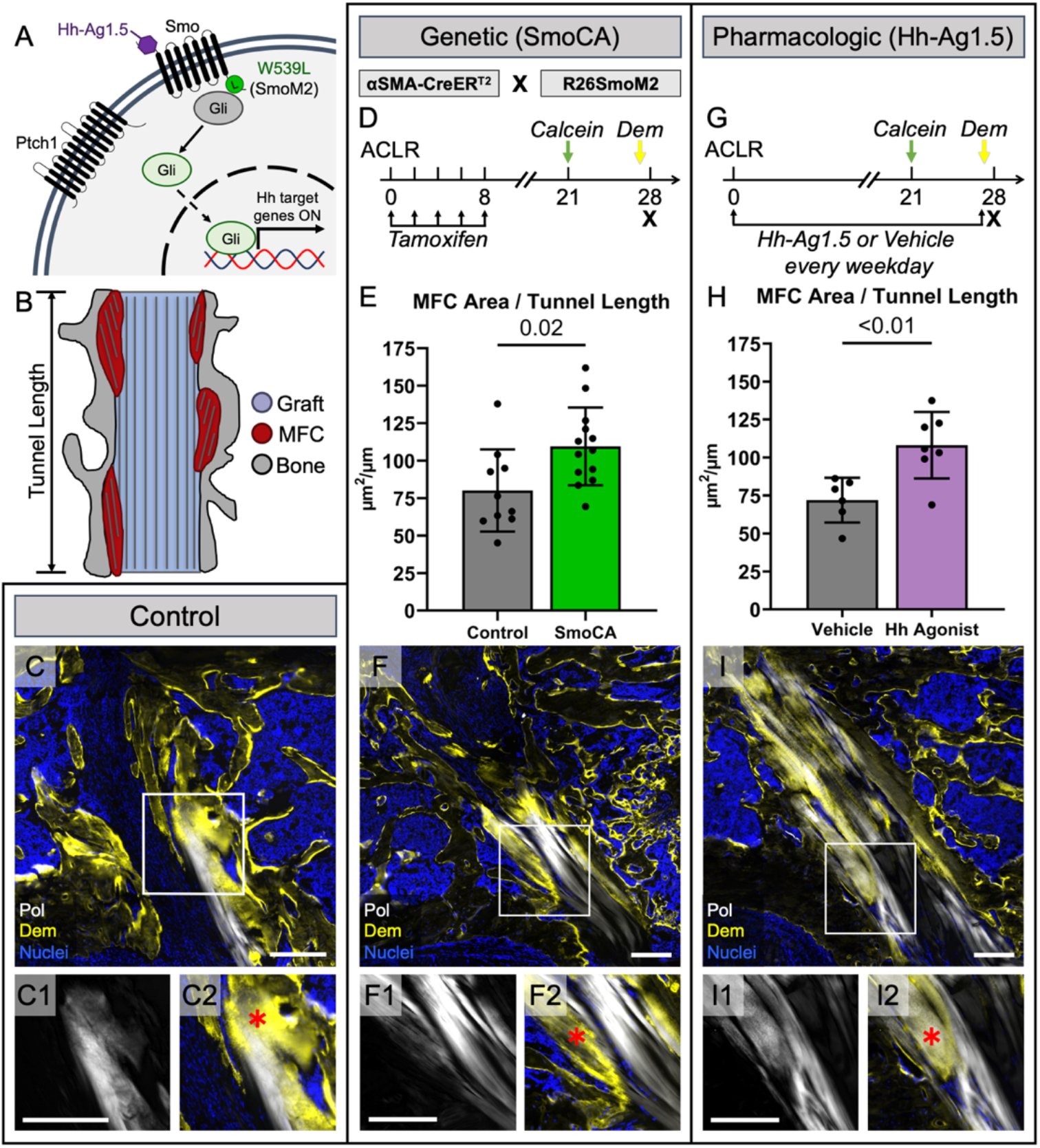
Hh signaling promoted zonal attachment formation following ACL reconstruction. Without Hh ligand binding, the Hh pathway (A) is inhibited by Ptch1 suppressing the activity of Smo and maintaining Gli proteins in their repressed form. Upon ligand binding to Ptch1, the Smo inhibition is removed and Smo can convert Gli proteins into their active state, leading to downstream Hh-target gene expression. Mineralized frozen sections were made at day 28 post-surgery and we quantified MFC area (B) by outlining areas where collagen fibers (white in C, C1, C2) displayed Dem mineral label (yellow in C, C2) and normalized the area by the tunnel length. We genetically modulated the Hh pathway by incorporating a mutant form of Smo, SmoM2, that leads to Ptch1 and ligand-independent activation of the pathway and crossed these mice with αSMACreERT2 mice (SmoCA) that target the cells participating in attachment formation (D). We gave tamoxifen injections in the first week post-surgery and mineral labels 7 and 1 day before sacrifice to label deposited mineral. SmoCA mice had increased MFC area 28 days post-surgery (E) (p = 0.02, n = 10-13/group). Representative sagittal section from SmoCA mice is shown in (F), with insets showing collagen structure (F1) and MFC (F2). We also gave systemic injections of the Smo agonist, Hh-Ag1.5, (A) every weekday for 28 days post-surgery (G), which increased MFC area (H) (p < 0.01, n = 6-7/group). A representative section from an agonist-treated mouse is shown in (I). Scale bars = 200μm. Red ✱ in C2, F2, and I2 represent MFC.

Since genetic overexpression of Smo had a positive effect on MFC formation 28 days post-surgery, we next determined whether a therapy using the small molecule Hh agonist (Hh Ag1.5), which binds to Smo regardless of Smo inhibition by Patched1 (Ptch1) (Fig. 2A), would have a similar effect on tunnel integration. To confirm the effectiveness of agonist injections on increasing Hh activity within cells participating in the repair response, we gave daily injections and measured expression of Gli1 and Ptch1 in regions adjacent to the bone tunnels on day 7. This dosing regimen yielded 16.2-and 11.7-fold increases in Gli1 and Ptch1, respectively (Fig. S2). After demonstrating effectiveness using this dose, we then gave injections every weekday for 28 days post-surgery (Fig. 2G) and compared the MFC formation to vehicle controls. Hh agonist-treated mice had 50% greater MFC area after 28 days compared to mice that received vehicle injections (p < 0.01, Fig. 2H, I), indicating that Hh signaling agonist delivery promotes tendon-to-bone integration.

### Up-regulating Hh signaling improves tendon-to-bone integration strength following ACL reconstruction

To directly test whether the increased MFC in Hh-targeted mice resulted in higher integration strength, we performed tunnel pullout tests on SmoCA and Hh agonist-treated mice as previously described^18^. The knee was disarticulated and the femur and tibia were separated. Femurs were used for pullout tests because the tendon knot around the tibial washer was not strong enough to consistently withstand the forces required in the pullout test. While SmoCA mice only displayed an increasing trend (p = 0.25, Fig. 3A, left), Hh agonist-treated mice yielded 48% greater pullout strength at 28 days post-surgery compared to vehicle controls (p = 0.03, Fig. 3A, right). Interestingly, the failure location in the SmoCA mice, and their Cre-negative littermates, was shifted towards the graft region outside the tunnel adjacent to the washer compared to the intra-tunnel failures in the agonist-treated and vehicle control mice (Fig. 3B). This shift in failure location may have negatively impacted the results of the SmoCA study. Nonetheless, these results indicated that the increased MFC formation in the Hh-treated mice resulted in improved tunnel integration strength.

**Figure 3.**
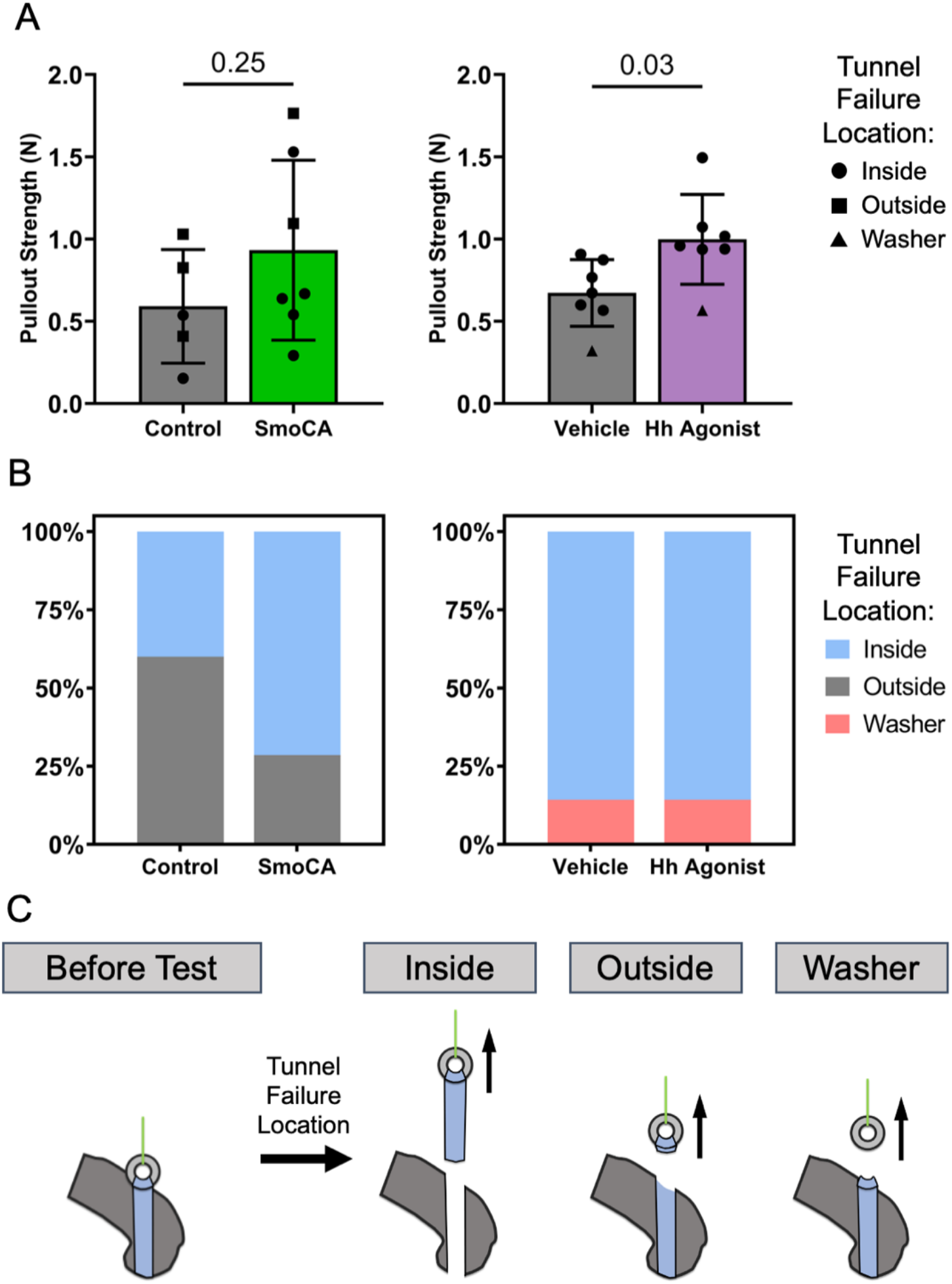
Hh signaling improves tendon-to-bone integration strength following ACL reconstruction. Femurs were dissected of soft tissue and mounted in PMMA. Fishing line was hooked through the femoral washer on one end and a grip on a linear actuator on the other end. The washer was pulled linearly until failure and the maximum failure force was recorded. SmoCA mice (A, left) and Hh agonist-treated mice (A, right) had greater pullout strengths compared to their respective controls (p = 0.25, n = 5-7/group for SmoCA, p = 0.03, n = 7/group for Hh agonist). Failure locations for each sample were recorded and were categorized into i) inside tunnel, ii) outside tunnel, and iii) washer failures. Failure locations for each sample are denoted in A with shapes and the percentage distributions for failure locations are plotted in B for SmoCA (B, left) and Hh agonist-treated mice (B, right). Schematics to help visualize failure locations are shown in C.

### Hh signaling promotes proliferation of the amplifying progenitor pool

Since our results indicated that Hh stimulation increased tendon-to-bone integration, both genetically and pharmacologically, we next sought to determine the mechanism by which Hh signaling improved this process. Given that the Hh pathway was previously shown to promote proliferation of bone marrow-derived mesenchymal stromal cells^42^, in addition to several other cell types^43–49^, we determined the effect of agonist treatment on proliferation of the expanding bMSC progenitor pool. By increasing expansion of this pool, there would be more cells capable of forming attachments, which could contribute to the increased MFC formation at day 28 (Fig. 2). Therefore, we delivered daily injections of Hh-Ag1.5 or vehicle in addition to daily injections of EdU for one week post-surgery and assessed the percentage of EdU positive cells within the expanding bone marrow adjacent to the bone tunnels at 7 days post-surgery (Fig. 4A). Since these regions contained newly formed woven bone that harbored EdU+ cells (Fig. 4A1), we excluded the woven bone from our analysis (Fig. 4A2). We found that agonist treatment yielded a 52% increase in cell proliferation (p = 0.03, Fig. 4B) and a 18% increase in cell density in the expanding marrow (p < 0.01, Fig. 4C) compared to vehicle controls, suggesting that pharmacologic hedgehog stimulation results in a larger pool of cells capable of forming tendon-to-bone attachments.

**Figure 4.**
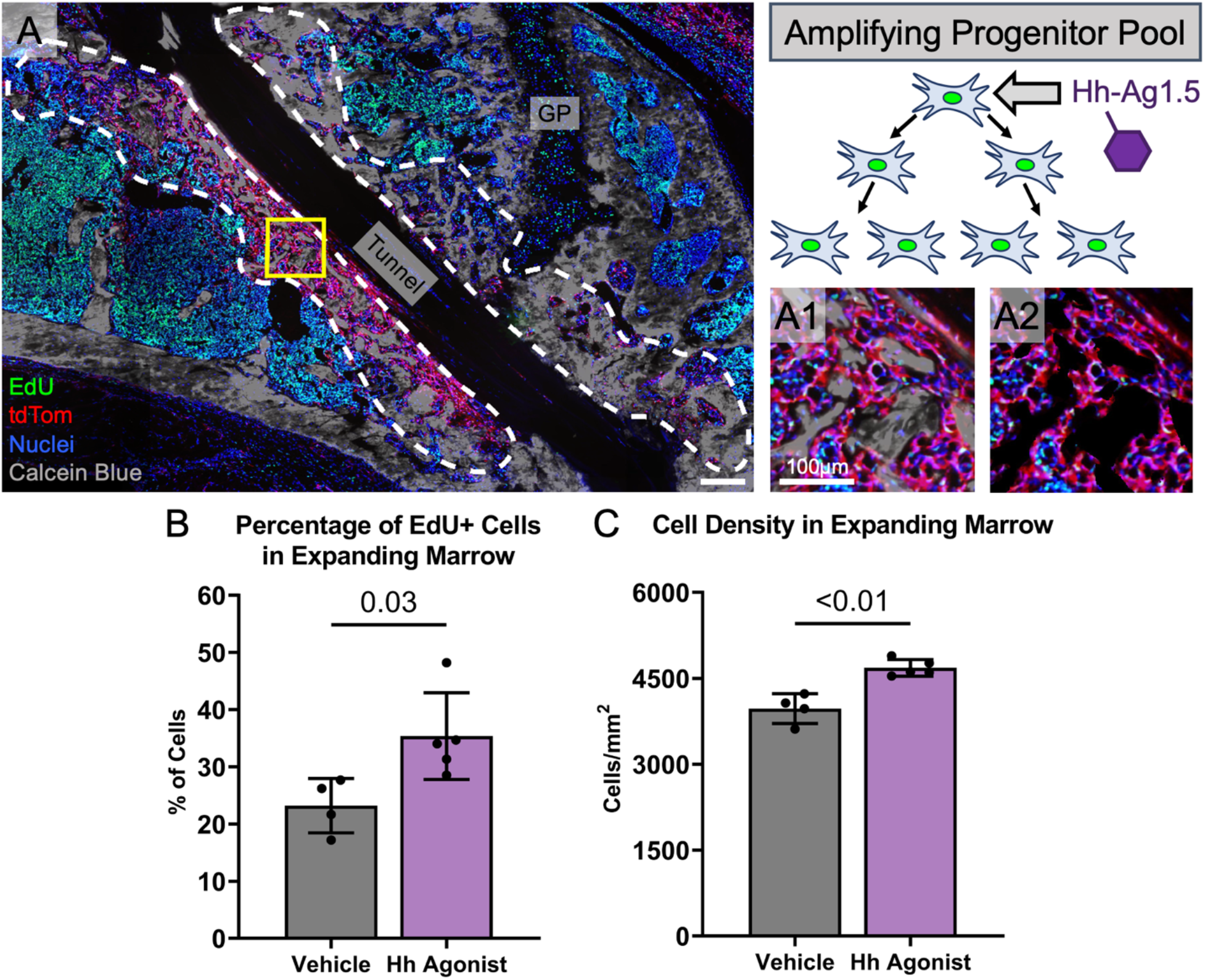
Hh stimulation in the first week post-surgery increased proliferation of the amplifying progenitor pool. αSMACreERT2;R26R-tdTomato mice that received Hh-Ag1.5 and EdU for 6 days post-surgery were assessed at 7 days post-surgery to determine the effect of Hh agonist treatment in the first week of surgery on proliferation of the amplifying bMSC progenitor pool. The area around the tunnel where tdTomato cells were present in the expanding marrow was used for analysis (dashed white outline in A). Because newly formed woven bone was present in this region at 7 days post-surgery (gray in A1), the EdU cell count contributions from the bone areas were removed from the final EdU+ cell counts (A2). Hh agonist treatment increased the proliferation of cells in the expanding marrow (B) (p = 0.03, n = 4-5/group) which resulted in an increased cell density in this region (C) (p < 0.01). GP = growth plate. Scale bar in A = 200μm.

### Hh stimulation improves tunnel integration as a result of more attachments along the tunnel length and increased mineral apposition within each attachment

Since Hh agonist treatment increased proliferation of the bMSC progenitor pool that gives rise to zonal attachments, there are conceivably more cells available to infiltrate the graft, differentiate into mineralizing fibrochondrocytes, and create zonal attachment sites, which would ultimately lead to more attachment sites along the tunnel length (Fig. 5A). To test this mechanism, we took the sections used for MFC area analysis and projected the areas of MFC horizontally to the edges of the tunnel (Fig. 5B) to obtain the length of the tunnel containing MFC (M_L_), which was then normalized by the total length of the tunnel (T_L_) (Fig. 5B1) to calculate the percentage of the tunnel length containing MFC. We found that SmoCA mice had 17% greater percentage of tunnel length containing MFC compared to controls (p < 0.01, Fig. 5C). Additionally, Hh agonist-treated mice had 23% greater percentage of tunnel length containing MFC compared to vehicle controls (p = 0.02, Fig. 5D). These results indicate that Hh stimulation increased the number of attachment sites formed, which may be attributed to the increased proliferation of the bMSC progenitor pool.

**Figure 5.**
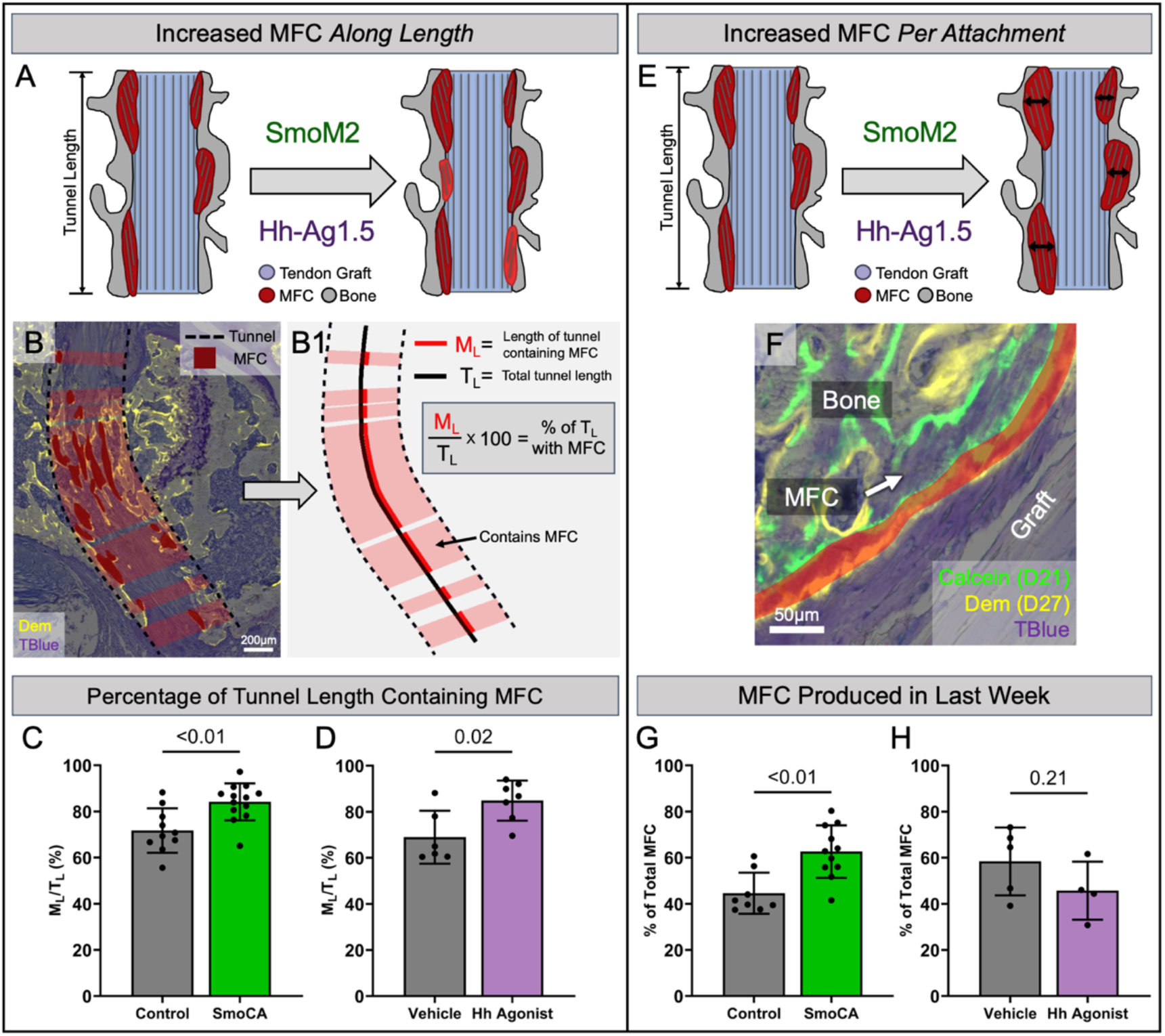
Mineralized fibrocartilaginous attachments increased after Hh stimulation as a result of more attachments along the tunnel length and increased mineral apposition within each attachment. Mice that were used for MFC area analysis from SmoCA and Hh agonist-treated mice were analyzed to determine the mechanism for increased MFC area: more attachments along the tunnel length (A) or more MFC produced in the last week as a result of more fibrocartilaginous differentiation (E). The MFC area regions used for MFC area/tunnel length quantification (dark red regions in B) were extended horizontally to the edges of the tunnel (dashed black lines in B) to obtain the length of the tunnel containing MFC (M_L_) which was then normalized by the total length of the tunnel (T_L_) to get the percentage of the tunnel length containing MFC (B1). The percentage of the tunnel length containing MFC was increased in the SmoCA mice (C) (p < 0.01, n = 10-13/group) and after Hh agonist treatment (D) (p = 0.02, n = 6-7/group) 28 days post-surgery. To determine if the mechanism for greater MFC area was due to more MFC production in the last week post-surgery, SmoCA and Hh agonist-treated mice received a calcein label at day 21 and a demeclocycline label at day 27 to label mineral deposited in the last week, and the MFC area between the two labels was measured (red in F). SmoCA mice had greater percentage of MFC produced in the last week compared to controls (G) (p < 0.01, n = 8-11/group), while Hh agonist-treatment did not produce a difference in MFC produced in the last week (H) (p = 0.21, n = 4-5/group). Although the same samples were used for quantifying both the percentage of the tunnel length containing MFC and MFC produced in the last week, the lower sample size for the latter measurement was due to missing calcein mineral labels.

Another potential mechanism that could explain the increased MFC area is an increase in mineral apposition by fibrochondrocytes in each attachment (Fig. 5E), which would indicate that Hh signaling promoted fibrocartilage maturation similar to its role in enthesis growth and development^20,23,25^. To test this mechanism, we measured the MFC produced during the last week by subtracting the MFC area up to day 21 (defined by the calcein label given on day 21) from the total MFC area (defined by the demeclocycline label, see Fig. 2) in SmoCA and Hh agonist-treated mice (red shaded region in Fig. 5F). Similar to total MFC (Fig. 2), SmoCA mice had 40% greater percentage of MFC produced in the last week post-surgery compared to controls (p < 0.01, Fig. 5G) while Hh agonist treatment did not significantly affect the amount of MFC produced in the last week compared to vehicle-treated mice (p = 0.21, Fig. 5H, breakdown of MFC produced before and after day 21 can be found in Fig. S3). In addition to the proliferation results in Fig. 4, these findings suggest that Hh signaling has a biphasic role in tendon-to-bone attachment formation in this model.

### Hh stimulation did not affect new bone formation around the tunnels

Since the activated αSMA-expressing progenitor pool gives rise to new bone adjacent to the tunnels, in addition to the MFC within attachments in the tunnels, and Hh signaling promotes osteogenesis^50–60^, we tested whether Hh promoted new bone formation adjacent to the bone tunnels in our model. To determine whether increased proliferation within the expanding marrow after Hh agonist treatment resulted in increased production of newly formed woven bone around the tunnels, we quantified the Calcein Blue-stained mineral in the same region around the grafts that we used for EdU analysis and found no difference in the amount of bone in this region nor the total expanding marrow area (Fig. S4). Although Hh stimulation did not increase bone formation after 1 week, we next sought to determine whether sustained Hh stimulation would lead to more bone around the tunnels at 28 days-post surgery. We used OsteoMeasure to quantify histomorphometric parameters approximately 400µm from the tunnel interface in SmoCA and Hh agonist-treated mice 28 days post-surgery (Fig. S5). All parameters, except MAR of the SmoCA mice (p < 0.03, Fig. S5D), were not significantly different in Hh-stimulated mice compared to controls, indicating that while Hh stimulation improves MFC formation it had minimal effect on bone formation in this model.

## DISCUSSION

Tendon-to-bone integration following ACL reconstruction occurs through a coordinated series of events, initiating with amplification of the bMSC progenitor pool, infiltration of these cells into the graft, and culminating in the formation of zonal fibrocartilaginous tendon-to-bone attachments. This study demonstrates that Hh signaling has a biphasic role in this coordinated process by positively regulating the proliferation of the expanding progenitor pool and also promoting the production of MFC in the zonal tendon-to-bone attachments in the bone tunnels (Fig. 6). While it has previously been established that the Hh pathway is active during tendon-to-bone repair^13,28,29^ and may have a positive role in enthesis healing^61^, we found that as the cells differentiate, the levels of Gli1, Ptch1, and Ihh expression increase as well, indicating that Hh activity becomes more prominent during the later stages of zonal attachment formation, likely within Col2a1 and Col10a1-expressing mineralized fibrochondrocytes (Fig. 1D). Unlike the previously used ScxCre;Smo^flox/flox^ conditional knockout model that has developmental defects^20,23,25^ that may influence the tendon-to-bone healing response^61^, we utilized inducible αSMACreERT2 mice to activate hedgehog signaling in cells after they initiate the repair response. This genetic approach yielded increased formation of zonal tendon-to-bone attachments (Fig. 2D-F), indicating that activating Hh signaling within the αSMA-expressing progenitor pool will improve their ability to form MFC within these zonal attachments. We also utilized the small molecule agonist, Hh Ag-1.5, to stimulate the pathway therapeutically, which produced similar improvements in tendon-to-bone integration (Fig. 2G-I). Taken together, these results indicate that the Hh pathway is a potent regulator of the tendon-to-bone integration process.

**Figure 6.**
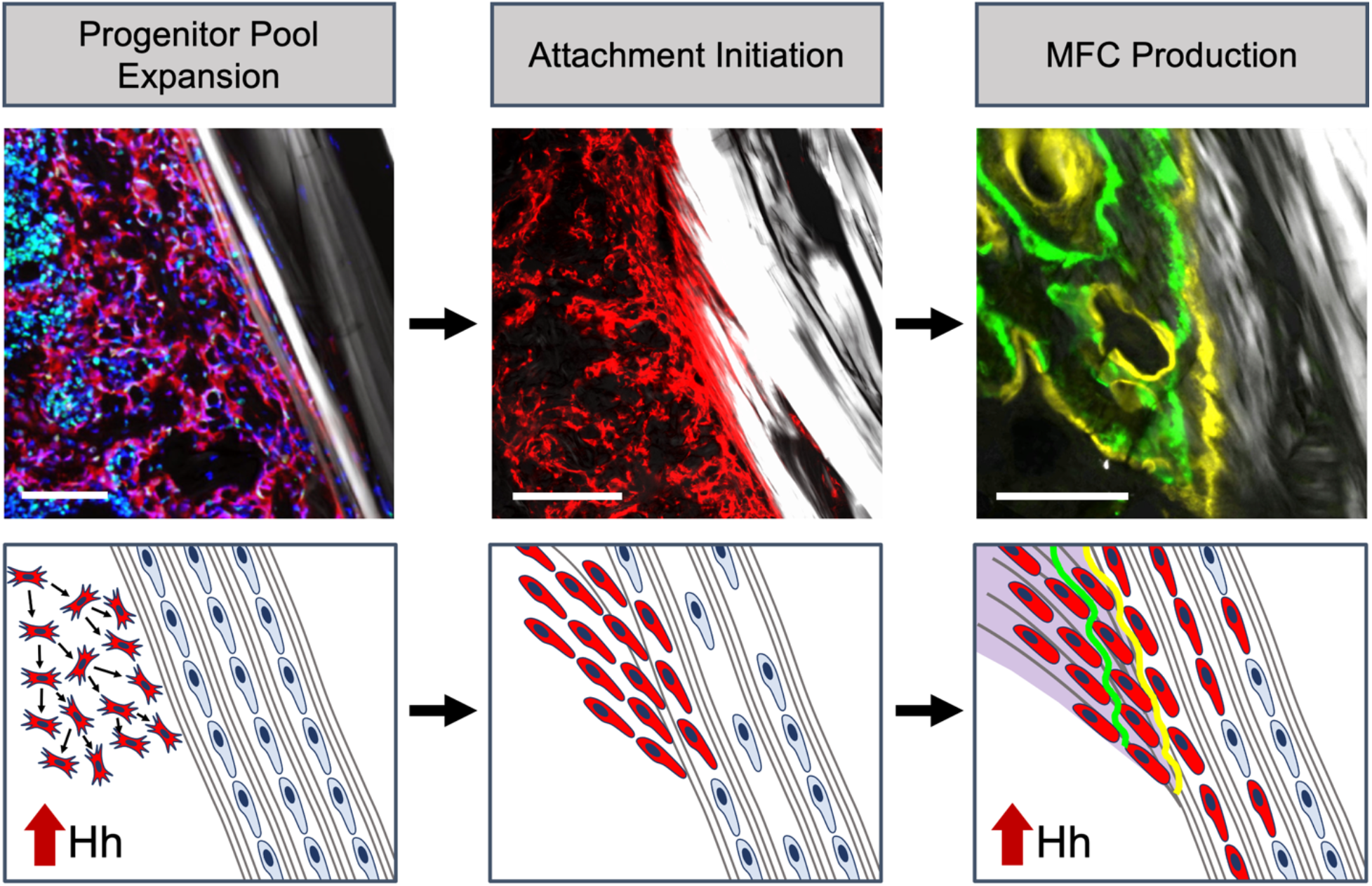
Hh signaling has a biphasic role in the tendon-to-bone integration following ACL reconstruction. The tunnel integration process begins with bMSC progenitor pool expansion in response to the drilling of the bone tunnels. We found that hedgehog signaling plays a role in promoting expansion of this progenitor pool, leading to a greater population of cells capable of initiating the graft and creating attachments. These cells then infiltrate the tendon graft to initiate the attachments, which coincides with cell death of native tenocytes within the graft. Finally, the infiltrating cells synthesize proteoglycan-rich extracellular matrix (purple) and mineralize this region in an appositional manner, leading to the creation of distinct zones of bone, mineralized fibrocartilage, unmineralized fibrocartilage, and tendon midsubstance.

Given that Hh signaling has a positive role in cell proliferation^42–49^, fibrocartilage differentiation ^20,23–27,62^, and osteogenesis ^50–60^, which are all stages of the tendon-to-bone integration process, we further investigated the influence of Hh signaling on these specific stages. These steps of the repair process are not independent, however. For instance, if there is a greater expansion of the bMSC progenitor pool, there will be a greater number of cells capable of producing a zonal attachment even if fibrocartilage differentiation is not affected. Interestingly, we found that Hh agonist delivery did indeed promote cell proliferation within the expanding marrow (Fig. 4), which is the source of cells that form the zonal attachments. Additionally, a larger proportion of the tunnel length contained MFC following Hh stimulation (Fig. 5A-D), suggesting that this proliferation effect led to an increase in the number of tendon-to-bone attachment sites along the tunnel length. On the other hand, if Hh signaling promoted fibrocartilage differentiation, we would expect there to be continued mineral apposition within each attachment site. In fact, we found higher production of MFC at later stages of tunnel integration in the SmoCA mice compared to controls (Fig. 5G). Taken together, these data indicate that Hh signaling has a positive role in both cell proliferation and mineralized fibrocartilage production in the formation of zonal tendon-to-bone attachments following ACLR, leading to improved tunnel integration strength (Fig. 3).

While genetic and pharmacologic Hh stimulation both increased MFC area (Fig. 2) and the percentage of the tunnel length containing MFC (Fig. 5A-D), only the SmoCA mice yielded an increase in the MFC produced in the last week (as a percentage of total MFC) compared to their respective control mice (Fig. 5G). When breaking down the MFC production to before and after day 21 (Fig. S3), it appears that SmoCA and agonist treatments have differential MFC dynamics. As seen in Figs. 5G, 5H and S3C, SmoCA mice produced more MFC than controls at later stages of the integration process, whereas Hh agonist treatment led to more MFC production in the earlier stages (Fig. S3D). Why daily systemic injections of the Hh agonist did not continue to stimulate MFC production could be due to a number of reasons. First, while αSMA-expressing cells in the bMSC progenitor pool and their progeny had constitutive and unrestricted activity of SmoM2 in the SmoCA mice, the agonist treated mice relied on continued delivery of the agonist to the tunnel space. The bone tunnels change drastically during the tunnel integration process as the attachments mature with more accumulated MFC. As this maturation occurs, diffusion of the agonist to these attachment sites is likely reduced, lowering the effectiveness of the agonist delivery. Additionally, Hh Ag-1.5 becomes less effective on cells at high concentrations^63^ suggesting that cells in these mice may have become desensitized to the agonist delivery at later stages of repair process. Future studies will investigate different agonist treatment windows, agonist concentrations, tamoxifen dosing regimens, and Cre drivers to further establish whether Hh plays a more prominent role in the early stages of proliferation and lineage commitment to fibrochondrocytes in the attachments or later stages of mineralized fibrocartilage production.

Because Hh signaling promotes osteogenesis ^50–60^, we investigated new bone formation adjacent to the bone tunnels in this model. Interestingly, agonist delivery did not increase woven bone formation at 7 days (Fig. S4) and neither the SmoCA nor agonist-treated mice demonstrated altered bone formation at 28 days post-surgery (Fig. S5). While we previously found that αSMA-expressing cells in the expanding marrow gave rise to fewer osteocytes than mineralized fibrochondrocytes in this model^18^, which may help explain why the SmoCA mice did not affect bone formation, we also did not see an effect with agonist delivery. This is in contrast with previous work in fracture healing^50–53,64^. However, there are a number of differences between bone formation in this ACLR model compared to fracture healing. For one, fracture healing is driven by periosteal progenitor cells whereas the new bone formation in ACLR is mostly bMSC-derived. Secondly, many of the fracture healing models used to study Hh signaling go through endochondral ossification whereas the bone formation following ACLR is mostly intramembranous. Ultimately, our data indicate that Hh signaling plays a more prominent role in the formation of fibrocartilaginous attachment sites than new bone in our ACLR model. However, further investigation into osteogenesis in our model is warranted.

This study is not without limitations. The tunnel pullout mechanical test is impacted by soft tissue scar formation around the femoral washer. Removal of this scar tissue requires careful dissection to remove the scar without damaging the adjacent tendon graft. We record videos of each test to not only help identify failure location of each specimen, but to help exclude samples where too much scar tissue remains. Interestingly, the failure location shifted in the SmoCA study compared to the agonist study in both treated and control mice. Unfortunately, many of the samples in the SmoCA study failed outside the bone tunnel adjacent to the washer, which limits our ability to test the tendon-to-bone interface in the tunnel (Fig. 3B). The reason for this difference in failure location is unknown. Additionally, the prolonged agonist dose of daily injections over several weeks is likely not a therapeutically viable and clinically translatable treatment regimen, as Hh signaling is essential in the maintenance of multiple tissues^65,66^. In our study, a small number of mice receiving Hh-Ag1.5 over the course of 28 days developed hair loss and skin hypertrophy at the injection sites around the third week of injection. Interestingly, a subset of SmoCA mice displayed heterotopic ossification lesions in the fat pad in the knee and around the patellar tendon at 28 days post-surgery (Fig. S6), but these lesions did not have a noticeable effect on limb ambulation. These lesions were likely due to overexpression of Smo in αSMA-expressing cells in the healing tissue in these regions. Ultimately, localized agonist delivery within a specific therapeutic window will be essential for harnessing the therapeutic potential of the Hh pathway in enhancing tendon-to-bone repair.

An improved understanding of the signaling pathways that regulate zonal insertion formation in the adult are crucial to developing new therapies to improve repair outcomes. By genetically and pharmacologically stimulating the Hh pathway in our ACL reconstruction model, we demonstrated that the Hh pathway plays a prominent, biphasic role in zonal attachment formation in the bone tunnels. If the Hh pathway is harnessed therapeutically, it could result in a paradigm shift in the clinical treatment of debilitating tendon and ligament injuries by improving repair outcomes, expediting recovery times, and potentially attenuating chronic disease progression.

## AUTHOR CONTRUBUTIONS

TBK, KF, SKL, XJ, RM, MHZ, AFK, and NAD conceived and designed the study. TBK, KF, SKL, XJ, RM, MKE collected and assembled the data. TBK, KF, SKL, XJ, RM, MKE, AFK, and NAD analyzed and interpreted the data. TBK, KF, SKL, MHZ, AFK, and NAD drafted the article. All authors provided critical revisions and final approval of the article.

## ACKNOWLEDGEMENTS

This study was supported by NIH R01-AR076381, R21-AR078429, R00-AR067283, F31-AR079840 (support for TBK), and P30AR069619 (pilot grant award and core facilities) in addition to the McCabe Fund Pilot Award at the University of Pennsylvania.

## SUPPLEMENTAL FIGURES

**Supplementary Figure 1.**
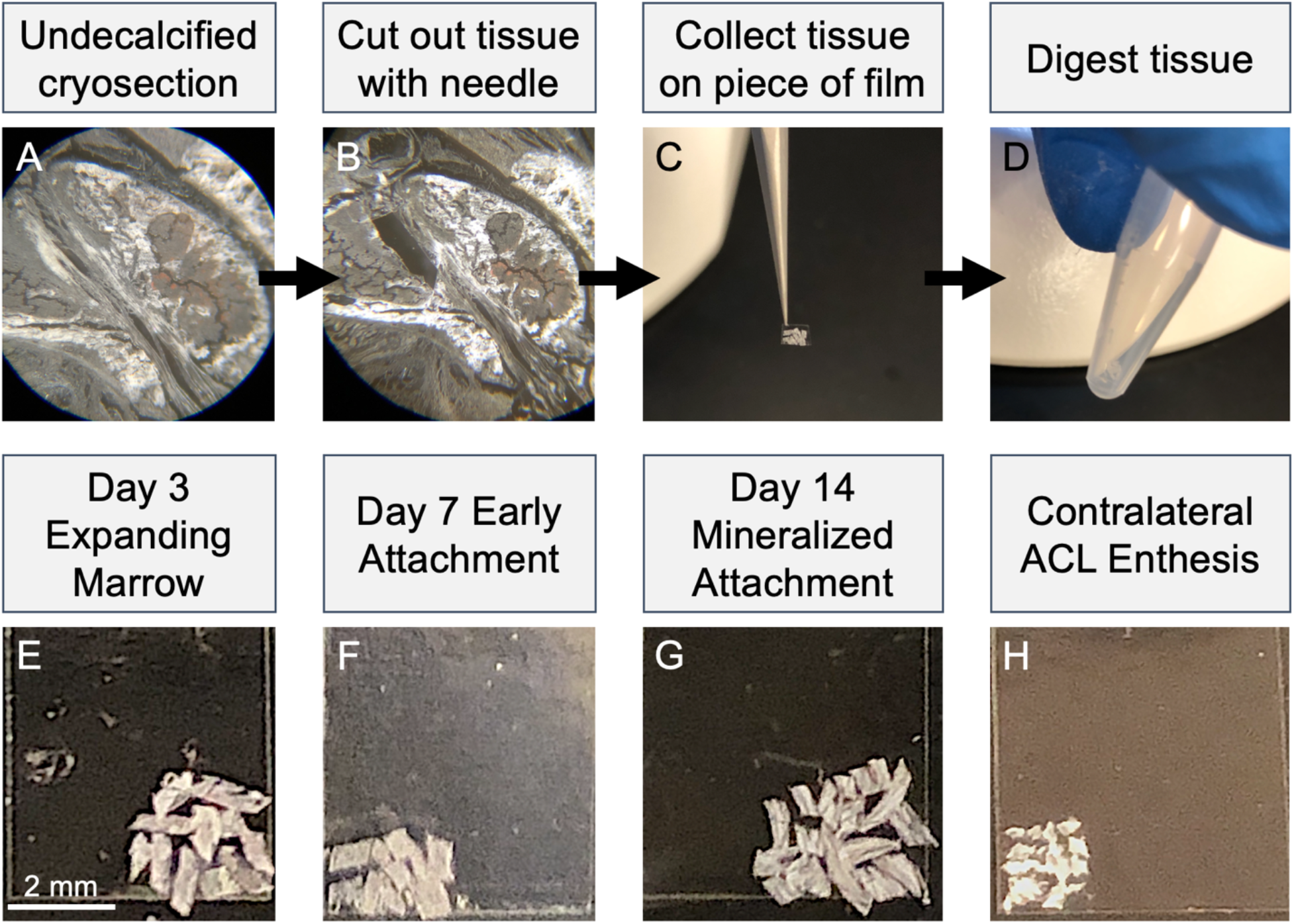
Microdissected cryosections for qPCR. Tape-stabilized 20µm thick cryosections were collected and kept frozen until tissue capture (A). Specific regions were cut from the sections under a dissection microscope using the beveled edge of a 27G needle (B), cutting through the tissue and cryofilm. The cryofilm-tissue pieces were collected on a separate, blank piece of cryofilm to pool tissue from multiple sections per sample (C). The cryofilm piece containing all dissected tissue from a sample was then transferred to a centrifuge tube for tissue digestion, RNA extraction, and further qPCR steps (D). Representative collected cryofilm-tissue pieces for the qPCR experiment performed in Figure 1D are shown for the day 3 expanding marrow (E), day 7 early attachment (F), day 14 mineralized attachment (G), and contralateral ACL enthesis (H).

**Supplementary Figure 2.**
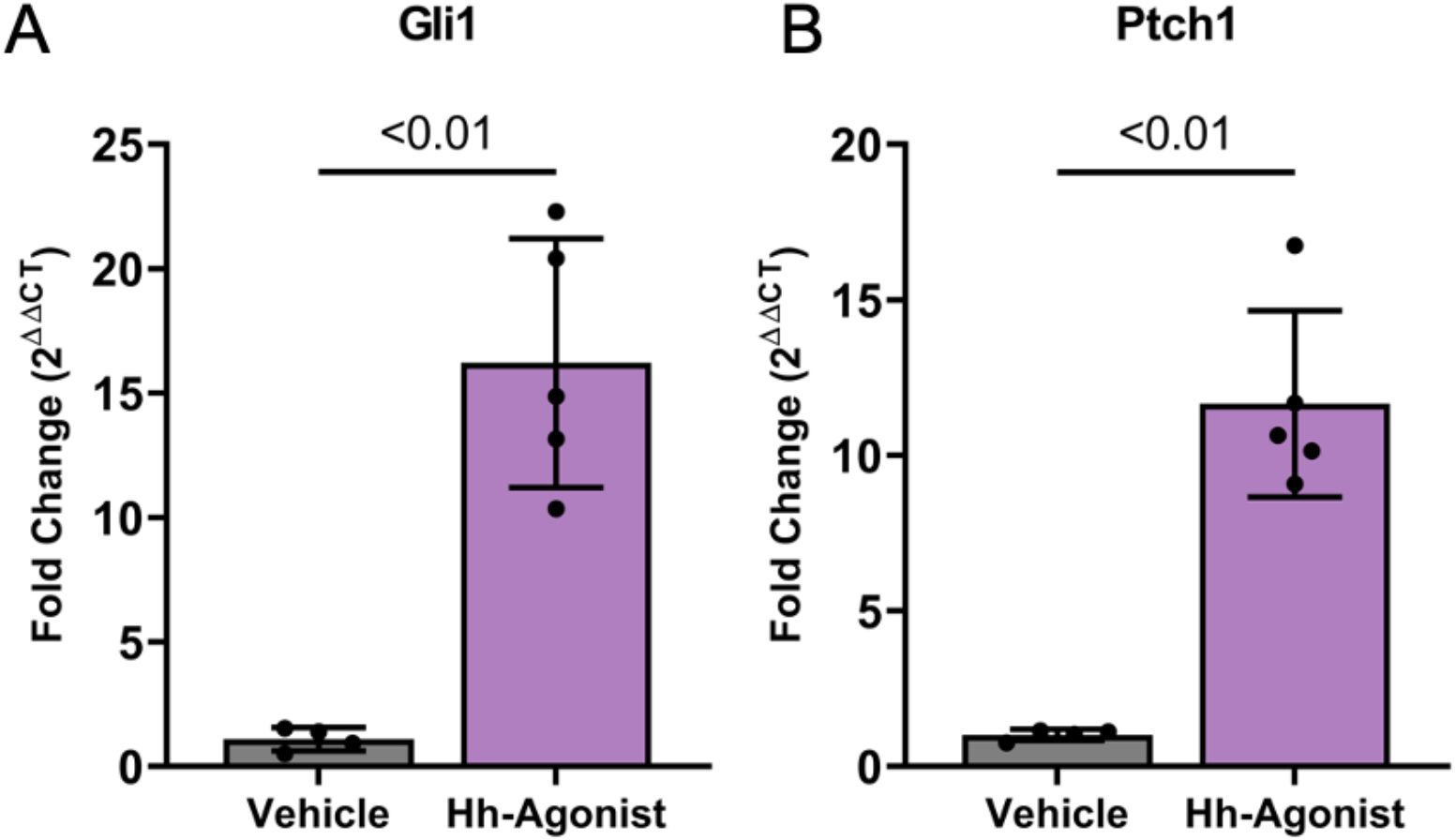
Hh agonist treatment increased Hh-related gene expression in the expanding marrow 7 days post-surgery. Cryofilm sections from samples used for proliferation analysis in Fig. 4 were subjected to microdissection of the expanding marrow around the tunnels and qPCR for Hh-related genes Gli1 (A) and Ptch1 (B). Both genes had significantly greater gene expression compared to vehicle controls (p < 0.01, n = 4-5/group).

**Supplementary Figure 3.**
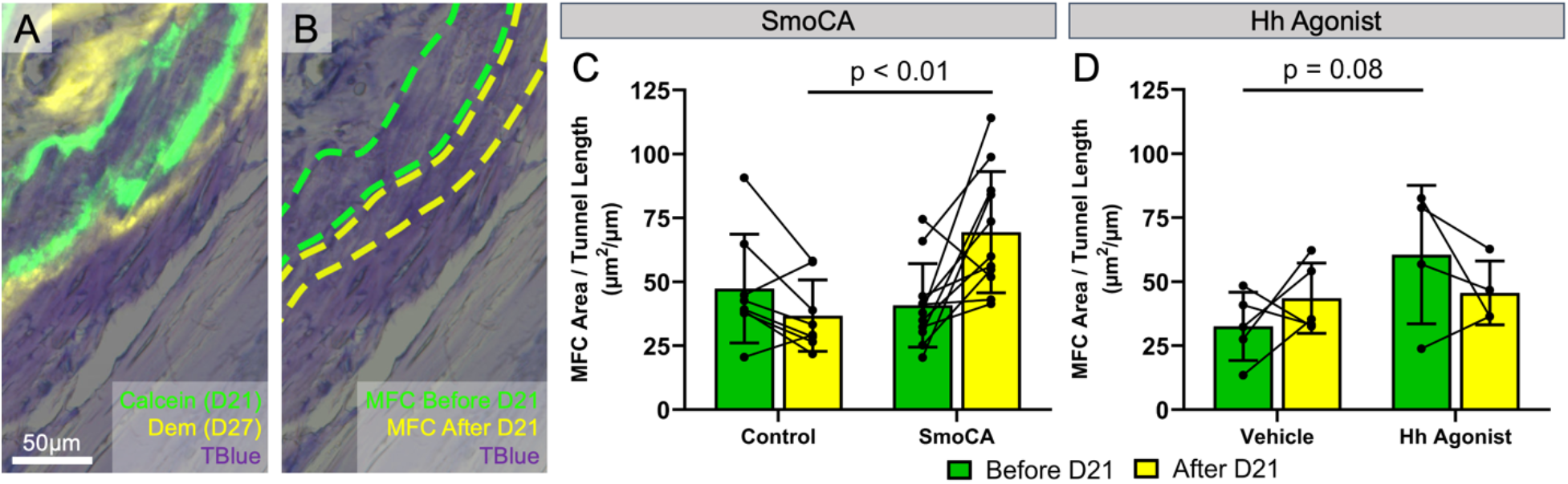
Temporal changes in MFC apposition between SmoCA and Hh Agonist treatment regimens. Calcein was delivered on day 21 (D21) and demeclocycline on day 27 (D27) post-surgery to determine the changing dynamics in MFC apposition as a function of healing time (A-B). Using the same features as those used to define total MFC in Fig. 2, we quantified MFC produced before and after D21 and noted them as “MFC before D21” (area enclosed by the dotted green line in B, green bars in C-D) and “MFC after D21” (area enclosed by the dotted yellow line in B, yellow bars in C-D). SmoCA mice (C) had no difference in MFC produced before D21 but had increased MFC produced after D21 (p < 0.01, n = 8-11/group), similar to the findings in Fig. 5G, where MFC after D21 was normalized to total MFC. Hh agonist-treated mice (D) had an increasing trend in the amount of MFC produced before D21 compared to vehicle controls (p = 0.08, n = 4-5/group) but no difference in MFC after D21.

**Supplementary Figure 4.**
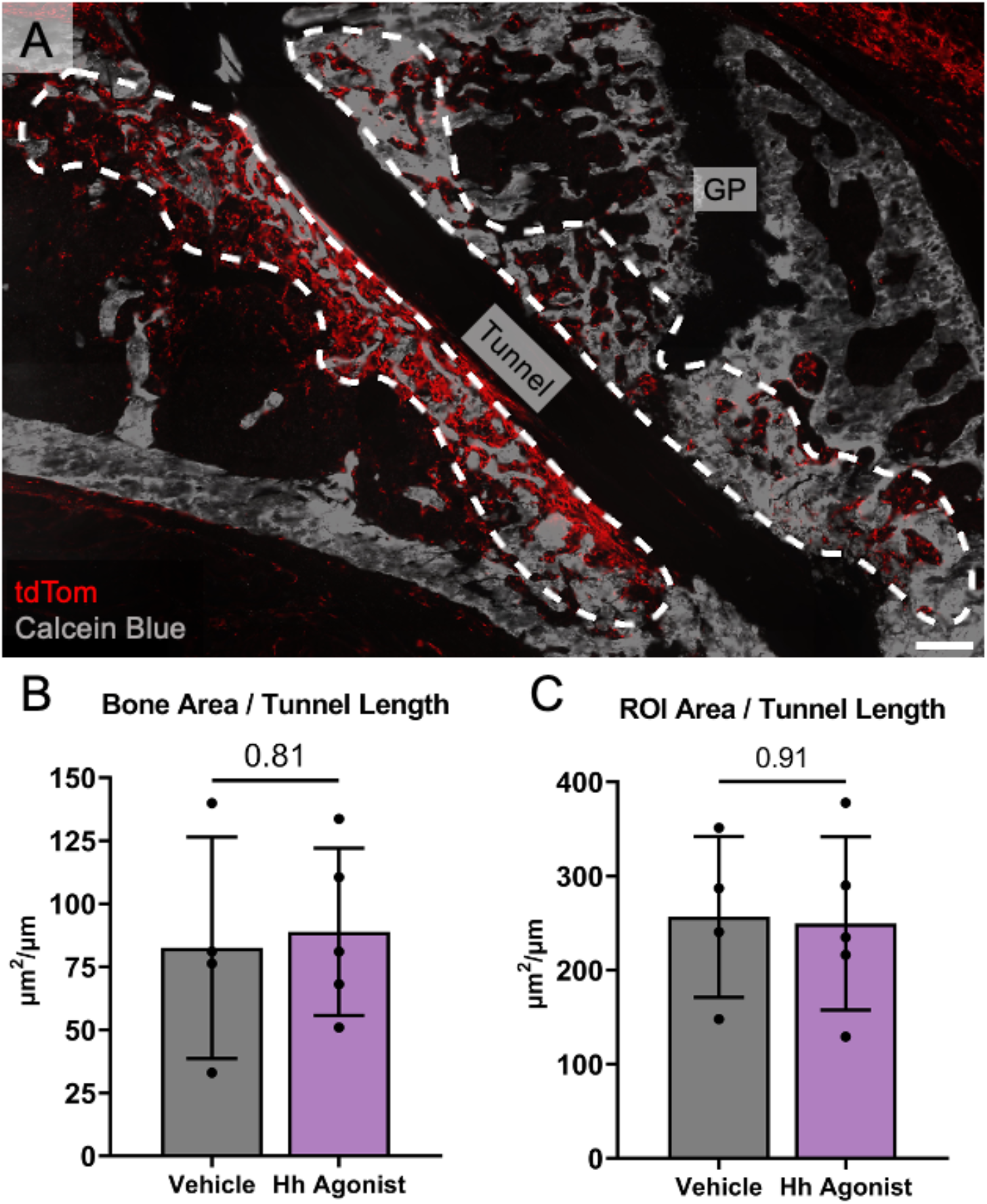
Hh agonist stimulation did not affect newly formed woven bone formation around the tunnel 7 days post-surgery. αSMACreERT2;R26R-tdTomato mice that received Hh-Ag1.5 and EdU for 6 days post-surgery were assessed at 7 days post-surgery to determine the effect of Hh agonist treatment in the first week of surgery on the early osteogenesis around the tunnel. The Calcein Blue stain for accumulated mineral (gray in A) was quantified in the expanding marrow around both sides of the tunnel containing tdTomato-positive cells (dashed outlines in A) (A). The ROI area for Calcein Blue quantification was the same as the ROI used for proliferation analysis in Fig. 4. Hh agonist treatment did not produce a difference in the total bone area in the expanding marrow normalized by the tunnel length (p = 0.81, n = 4-5/group) (B). To determine whether Hh agonist treatment increased the total area of expanding marrow around the tunnel, the total ROI area (dashed outlines in A) was calculated and normalized by the tunnel length. Hh agonist treatment did not increase the expanding marrow ROI area (p = 0.91, n = 4-5/group) (C). GP = growth plate. Scale bar = 200μm.

**Supplementary Figure 5.**
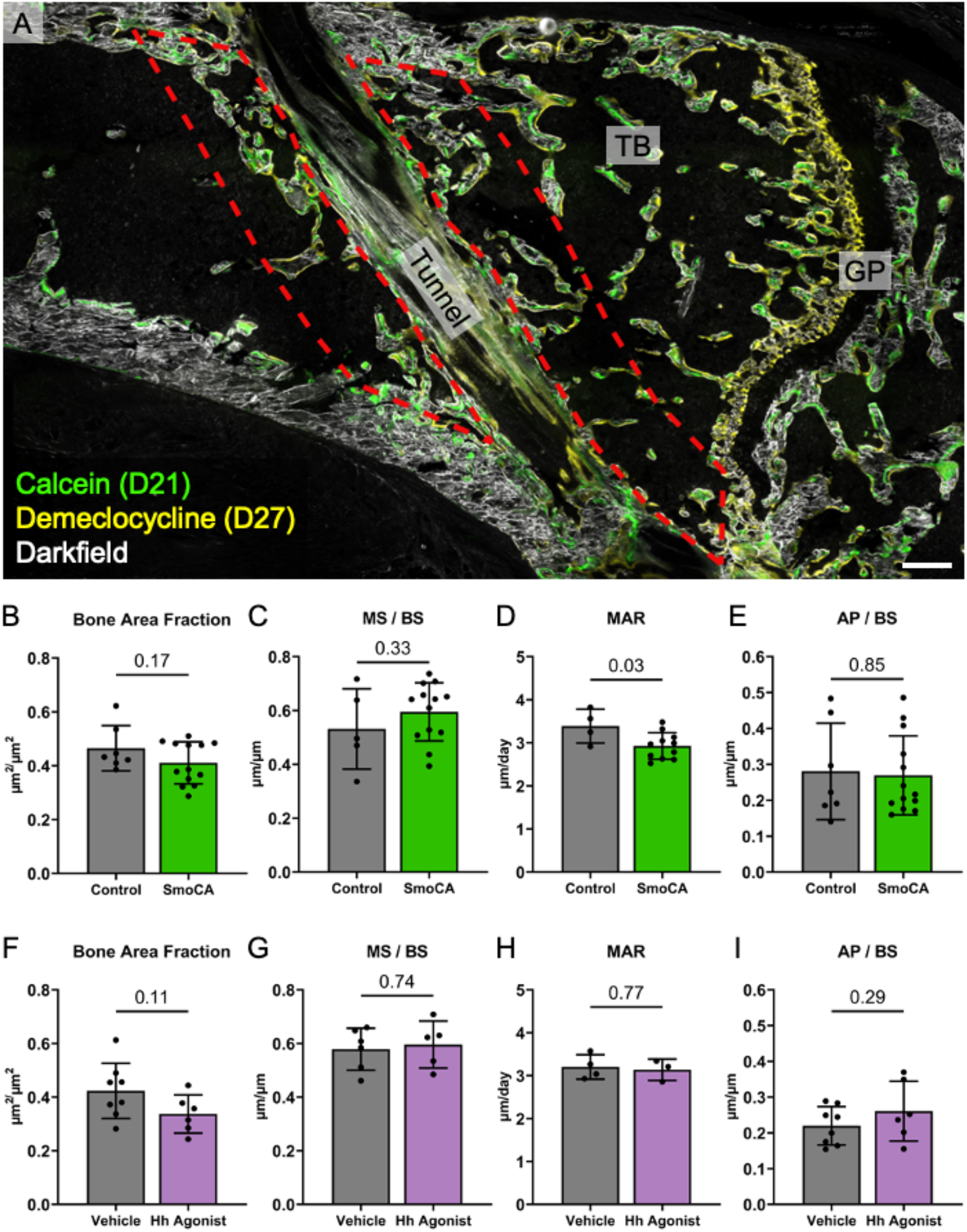
Hh stimulation does not increase bone formation around the tunnels after 28 days. Mice that were used for MFC area analysis from SmoCA and Hh agonist-treated mice were analyzed to determine whether Hh stimulation affects osteogenesis around the tunnels at 28 days post-surgery. Regions on both sides of the tunnel approximately 400μm away from the interface (red dashed outline in A) were used for bone parameter quantification in OsteoMeasure (A). Bone area fraction (B), mineralizing surface/bone surface (C), and alkaline phosphatase surface/bone surface (E) were not different in the SmoCA compared to controls. SmoCA mice had lower mineral apposition rate compared to controls (p < 0.03, n = 4-11/group) (D). BAF (F), MS/BS (G), MAR (H), and AP/BS (I) were not different in Hh agonist-treated mice compared to controls. GP = growth plate, TB = trabecular bone. Scale bar in A = 200μm.

**Supplementary Figure 6.**
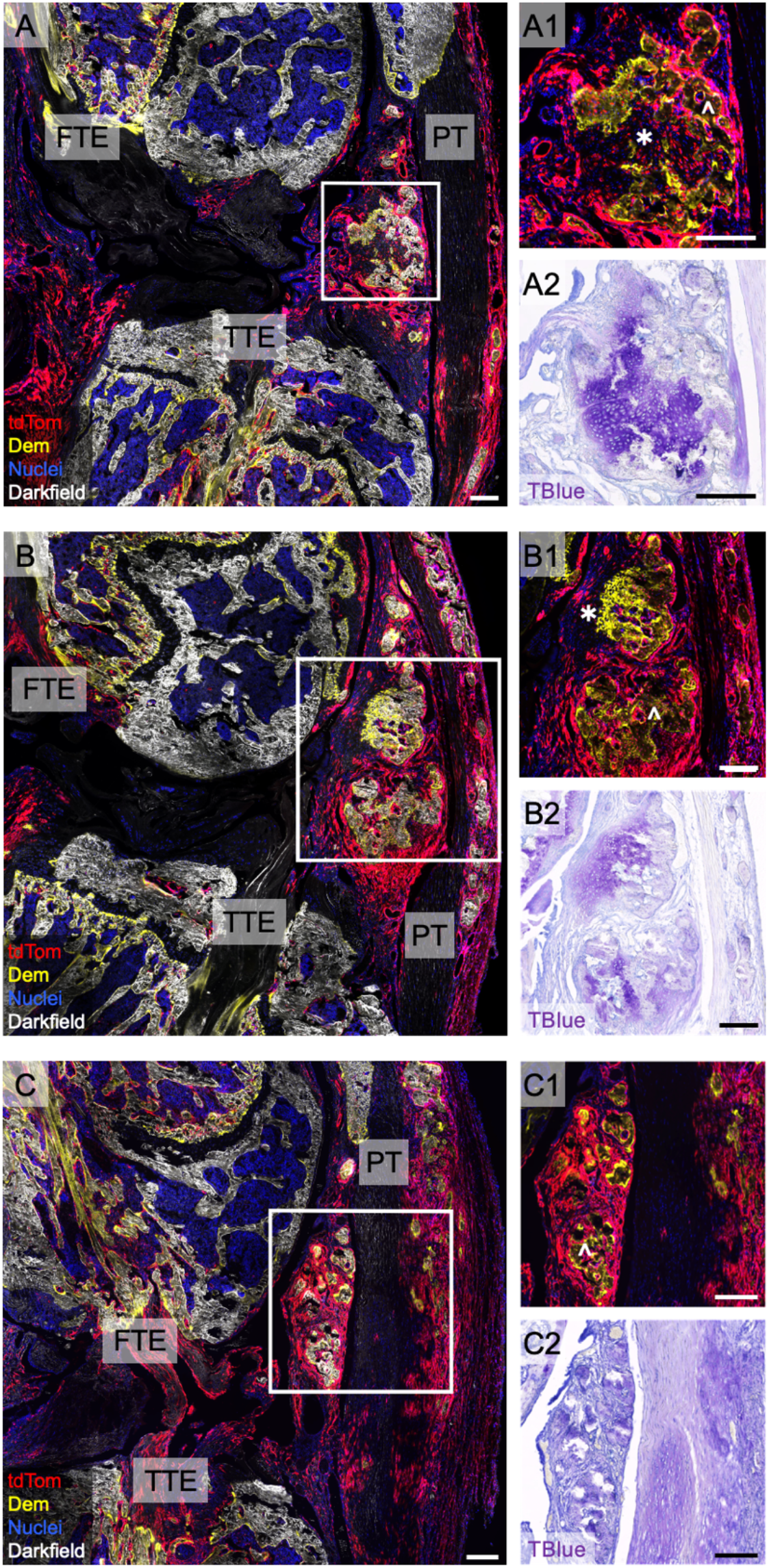
A subset of SmoCA mice displayed heterotopic ossification around the patellar tendon and fat pad 28 days post-surgery. Representative images from three SmoCA knee samples that had heterotopic ossification lesions in the fat pad and around the patellar tendon after 28 days (A, B, C). Some of these lesions (A, B) contained cartilaginous tissue (✱) with strong proteoglycan staining (A2, B2). Other parts of these legions contained bone tissue (^) (A1, B1, C1). FTE = femoral tunnel entrance, TTE = tibial tunnel entrance, PT = patellar tendon. Scale bars = 200μm.

